# PRISM : Peptide-specificity annotation of T-cell receptors with uncertainty quantification

**DOI:** 10.64898/2026.06.30.735715

**Authors:** Divya Lakshmi Venkatraman, Lydia Mok, Nicholas Rose, Andrew Robinson, Vanessa D Jonsson

## Abstract

Mapping T-cell receptor (TCR) sequences to their cognate peptide–major histocompatibility complex (pMHC) ligands underlies both basic immunology and T-cell target discovery, yet current models aimed at predicting TCR specificity are limited by sparse labels, viral-biased training data, and an inability to recognize receptors outside their training distribution. We present PRISM, an uncertainty-aware metric-learning framework for TCRβ sequence representation. PRISM embeds receptors into a peptide-organized latent space, returns top-k peptides by nearest-neighbor retrieval, and abstains on out-of-distribution receptors by modeling an intrinsic uncertainty that tracks annotation correctness. To offset the viral bias of public databases, PRISM augments training data with structure-guided synthetic receptors that diversify TCR sequences while preserving the energetics of the TCR–pMHC interface. Across a held-out set of 923 peptides and the independent IMMREP23 benchmark, PRISM matches or exceeds sequence-based models, with largest gains on rare epitopes. Finally, PRISM learns attention weights on TCR residues that concentrate on the CDR3β salt-bridge and hydrophobic contacts central to peptide recognition, linking PRISM’s positional focus to the biochemical properties of TCR–pMHC structures.

## Introduction

Adaptive immunity is mediated by a diverse T cell population whose receptors recognize peptide antigens presented by major histocompatibility complex (MHC) molecules^1^. High-throughput immune repertoire sequencing now resolves millions of T-cell receptor (TCR) sequences from a single sample, and identifies which TCR clones (T cells sharing an identical receptor sequence) are present and expanded, not which antigens they recognize ^2,3^. Experimental measurement of TCR specificity is feasible for only a small fraction of receptors in a repertoire, and sequence similarity does not reliably capture shared specificity. The receptor antigen relationship is degenerate, as near identical receptors recognize different antigens, while receptors with distinct sequences can converge on the same target^4,5^. Resolving antigen specificity from TCR sequence is thus a rate limiting step in mapping antigen specific immunity and identifying receptors for therapeutic engineering.

Existing databases including IEDB^6^, VDJdb^7^, McPAS-TCR^8^ and CEDAR^9^ aggregate experimentally characterized TCR–antigen pairs and make up the standard training data for specificity inference. Their coverage however is determined by which antigens are experimentally accessible, favoring immunodominant epitopes on common HLA alleles over the diverse restrictions that dominate real TCR repertoires. Viral epitopes are represented by thousands of receptors, whereas many tumor-associated, autoimmune, bacterial and rare pathogen specificities remain sparsely sampled or entirely absent^10^. This imbalance has two consequences for inference. First, a model estimated from such a distribution preferentially fits the densely sampled regions of recognition space and generalizes poorly to the underrepresented specificities of greatest translational interest. Second, for repertoire-scale application, the receptors requiring annotation in a clinical sample are those whose cognate antigens are absent from the reference data altogether, such that the predictions of greatest interest are precisely where the training data provide least support.

Computational approaches for TCR specificity inference broadly fall into two categories. Motif-and distance-based methods, including GLIPH2^11^, iSMART^12^, TCRMatch^13^ and tcrdist3^5^, group receptors by sequence similarity into candidate specificity sets and have proven useful for identifying antigen-associated sequence groups and disease-associated repertoire structure. However, they infer similarity among receptors rather than directly producing antigen-resolved predictions, and cannot capture receptors which share specificity without sharing sequence.

Supervised models instead learn TCR–peptide or TCR–peptide–MHC interactions directly from paired sequence data. Approaches such as NetTCR-2.0^14^, ERGO-II^15^, pMTnet^16^, PanPep^17^, MixTCRpred^18^ establish that specificity can be predicted from receptor and peptide inputs, yet recent independent benchmarks show that performance depends on negative sampling strategy^19^, epitope representation and whether the evaluated antigens are observed during training^17^. In particular, generalization to rare or unseen epitopes remains a major limitation in all models^20^.

A third approach reframes specificity as a problem of representation rather than classification. Embedding-based deep learning models such as TCR-BERT^21^ and SCEPTR^22^ learn TCR sequence representations transferable across downstream tasks such as specificity prediction, clustering, and repertoire analysis. Such representations aim to learn antigen-associated relationships from TCR sequence directly rather than by discrete class label. Metric-learning approaches are directly applicable here as they optimize the geometry of the embedding space itself^23^. These representations, however, lack an intrinsic measure of when a receptor lies beyond the reference, and most are not optimized for antigen-specificity retrieval under supervision—two capabilities on which reliable annotation at repertoire scale depends.

Addressing the coverage challenge would require expanding receptor diversity for underrepresented antigens as recent work indicates that the size and diversity of the training set, more than model architecture, govern generalization in TCR specificity prediction^24^. Modern inverse-folding and structure-guided protein design methods have shown that extensive sequence diversity can be generated while maintaining structural geometry and interface architecture, supporting the use of structural constraints to sample functionally plausible sequence variants^25^. This motivates an augmentation strategy for sparsely represented antigens with synthetic receptors generated under the biochemical and geometrical constraints imposed by the TCR–peptide–MHC molecular complex.

Here we present PRISM, a metric-learning framework that represents TCRβ sequences in an embedding organized by antigen specificity and reports, for each receptor, the reliability of its annotation. PRISM addresses the limitations of existing models through three mechanisms. PRISM embeds TCRβ sequences into a latent space using a supervised contrastive objective coupled to a ScaleFace loss, which places receptors of shared specificity in common neighborhoods while learning a per-receptor scale that quantifies local support within the same forward pass. To counter the reference’s bias toward immunodominant viral antigens, underrepresented antigens are enriched with structure-guided synthetic TCRs generated by `airr-gen`, which diversifies non-interface TCRβ complementary determining region (CDR) positions identified on predicted TCR-pMHC complexes. PRISM assigns antigen labels through embedding-based retrieval from neighboring experimentally characterized receptors, and provides the learned scale parameter as an intrinsic measure of uncertainty to complement annotation confidence. On a held-out set of 923 epitopes and the independent IMMREP23 benchmark, PRISM matches or exceeds sequence-based models, with the largest gains on the rare epitopes where reference data are thinnest, and assigns high uncertainty to out-of-distribution receptors rather than forcing them into known classes. Beyond annotation, PRISM outputs per-residue attention weights that correlate with the CDR3β residues that form specific bonds with the peptide, offering an interpretable view into features the embedding is built from.

## Results

### PRISM framework for antigen-specific TCR representation learning

We developed PRISM (Peptide–Receptor Inference via Supervised Metric-learning), a supervised metric-learning framework that learns antigen-informed representations from TCR beta chain sequences (Methods, Fig. 1). PRISM was trained on a harmonized dataset of antigen-specific TCRs assembled from public databases, and structure-guided synthetic data augmentation to expand receptor diversity for sparsely represented antigens (Fig. 1b). Because sequence similarity is an imperfect predictor of shared specificity (Supplementary Fig. 1 b,d), PRISM embeds TCRβ sequences into a latent space using a supervised contrastive-learning objective and a ScaleFace loss (Fig. 1c) which together draw receptors of shared specificity into common neighborhoods. Antigen assignments are subsequently obtained through distance-weighted K-Nearest Neighbors (k = 10) within the reference embedding space.

**Figure 1.**
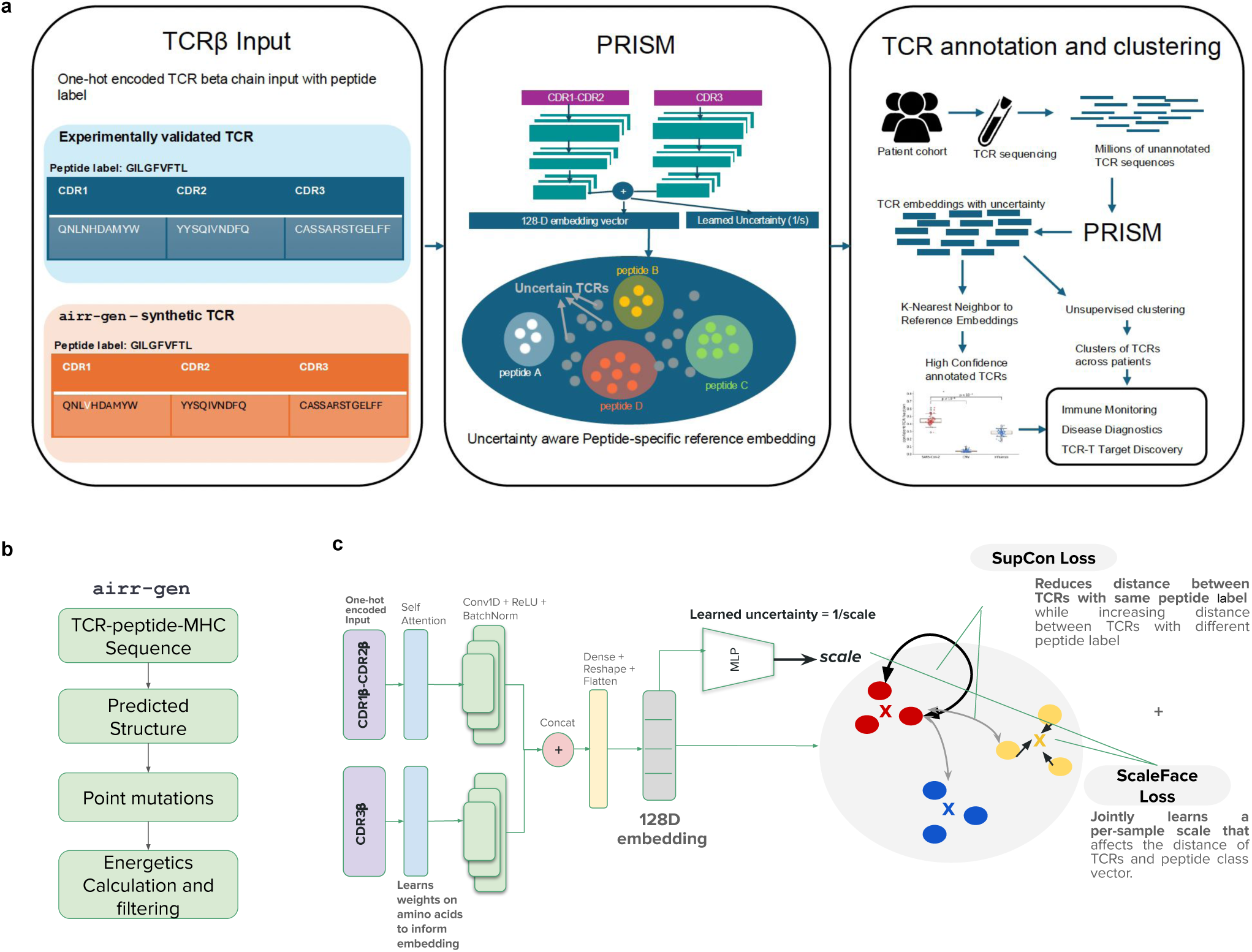
PRISM framework for antigen annotation of T-cell receptors. **(a) Overview of the PRISM framework.** (Left) TCRβ input is drawn from two sources: A curated set of experimentally validated TCR–antigen binding pairs, and airr-gen a structure-guided synthetic TCR augmentation pipeline (see b). Each TCR is represented by CDR1, CDR2, and CDR3 sequences of the beta chain, paired with a peptide label. (Center) PRISM learns a 128-dimensional embedding space in which TCRs targeting the same antigen (coloured clusters; peptides A–D) are brought together on a unit hypersphere. A per-TCR scale parameter *s* is learned jointly with the embedding; TCRs outside confident antigen-specific neighbourhoods (grey) receive high uncertainty (1/s) and are flagged for abstention. (Right) Downstream clinical workflow: unannotated TCR sequences from patient repertoires are embedded by PRISM and are annotated with K-nearest-neighbour retrieval against reference embeddings produces high-confidence antigen annotations for a subset of TCRs. Unsupervised clustering can be done to identify co-specific TCR groups across patients for immune monitoring, disease diagnostics, and immunotherapy discovery. **(b) airr-gen synthetic augmentation pipeline.** TCR–peptide–MHC sequence inputs are used to generate predicted structures, from which point mutations are introduced at non-interface CDR positions. Candidate sequences are filtered by computed binding energetics, producing synthetic TCRs that preserve antigen-contact residues while diversifying the surrounding CDR landscape to improve coverage of rare and tumour-associated antigen classes. **(c) PRISM model architecture and training objectives.** One-hot-encoded CDR1β–CDR2β and CDR3β sequences are processed through separate encoder branches. Branch representations are concatenated and projected to a 128-dimensional unit-hypersphere, yielding an embedding vector and a per-TCR scale parameter *s*. Attention weights are learned per amino acid position to focus the embedding on specificity-determining residues. The model is trained with two objectives: a supervised contrastive loss (SupCon) that minimises angular distance between TCRs sharing the same peptide label while maximising separation between TCRs with different labels, and a ScaleFace loss that jointly optimises *s* to modulate the effective classification margin per TCR. The learned uncertainty (1/s) is geometrically intrinsic to the embedding enabling confidence estimation and abstention on out-of-distribution receptors.

In addition to learning receptor embeddings, PRISM jointly learns a receptor-specific scale parameter through the ScaleFace loss that provides an intrinsic measure of uncertainty within the embedding space and outputs per residue attention weights that localize to the peptide-contacting CDR3β residues, giving a mechanistic view of which positions inform each representation. (Fig 1c). This framework enables simultaneous evaluation of antigen-specific organization, annotation performance and embedding confidence.

### PRISM learns antigen-specific TCR representations that support accurate specificity annotation

PRISM organizes TCRs by antigen specificity rather than sequence similarity. A t-SNE plot of the training embedding, corresponding to the 30 most frequently represented peptides, shows distinct peptide-associated neighborhoods (Fig. 2a): receptors recognizing the same antigen group together despite sequence diversity. The same separation holds on held-out test TCRs, where PRISM gives the only positive silhouette score against 10 peptide labels (0.105), while SCEPTR (−0.044), TCR-BERT (−0.080), and ESM2 (−0.157) do not separate peptides (Fig. 2b).

**Figure 2.**
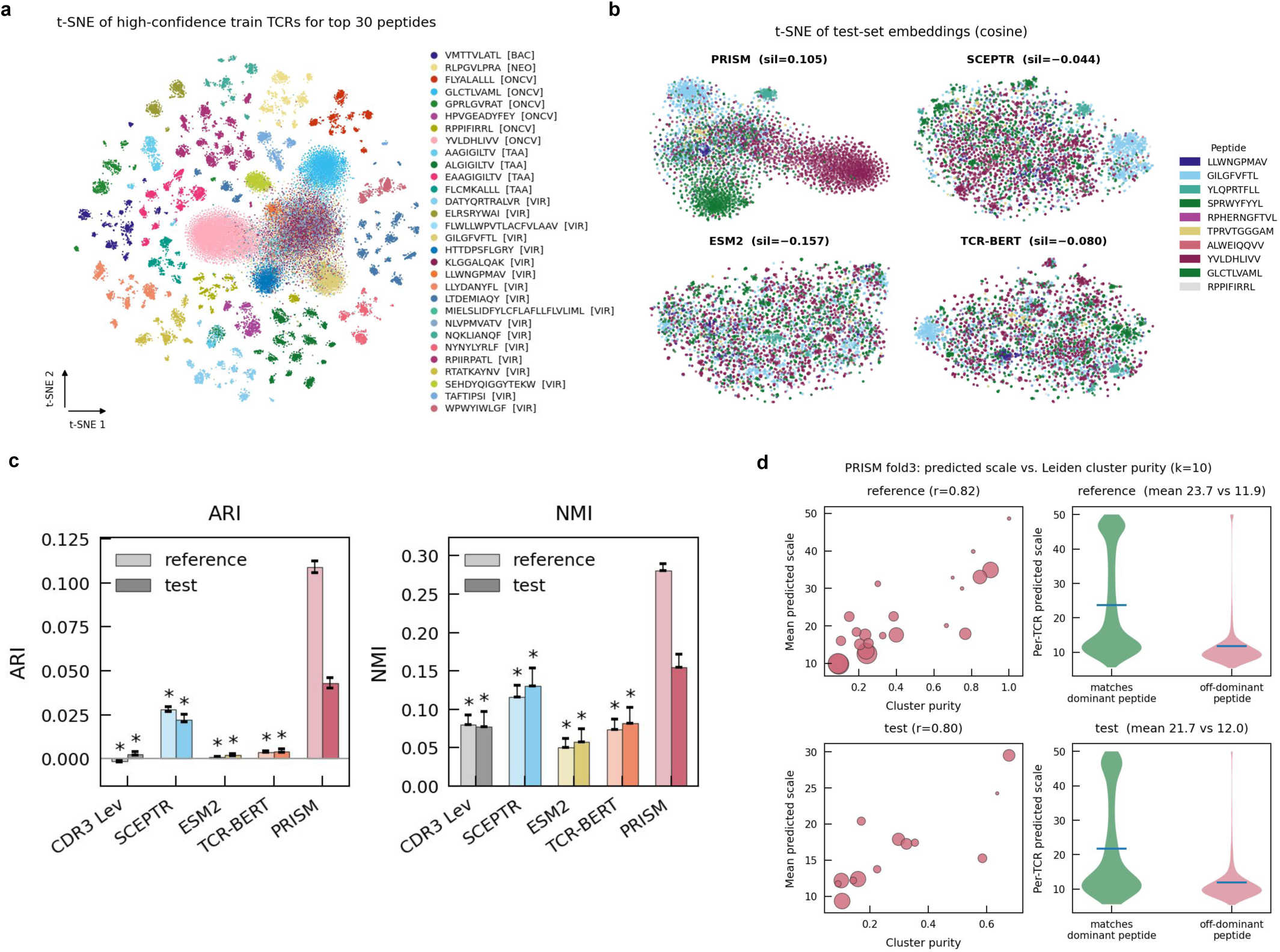
PRISM learns an antigen-resolved embedding space. **(a) t-SNE projection of trained PRISM embeddings.** High confidence (scale > 11) TCRs specific to the top-30 most abundant peptides in the training set are colored by peptide and annotated by antigen category (BAC, bacterial; NEO, neoantigen; ONCV, oncoviral; TAA, tumor-associated antigen; VIR, viral). **(b) t-SNE projections of held-out test-set embeddings for PRISM and three competitor models (SCEPTR, ESM2, TCR-BERT), colored by peptide specificity for the ten most represented test peptides.** Mean silhouette scores (cosine distance, computed over cells specific for these ten peptides) are shown above each panel; **(c) Adjusted Rand Index (ARI, left) and Normalized Mutual Information (NMI, right)** between Leiden clustering (k=10 nearest-neighbor graph, resolution=0.8) of model embeddings and ground-truth peptide labels, for CDR3 Levenshtein distance, SCEPTR, ESM2, TCR-BERT, and PRISM. Light bars, reference (training) set; dark bars, held-out test set. Error bars denote [95% CI across bootstraps per split]. Asterisks indicate significant difference from PRISM (* p < 0.05). **(d) Relationship between PRISM’s predicted uncertainty (scale parameter) and Leiden cluster quality** (k=10) for the reference (top row) and test (bottom row) sets. Left, per-cluster mean predicted scale versus cluster purity (fraction of cluster members sharing the cluster’s dominant peptide); point size proportional to cluster size (√n), Pearson r shown above each panel. Right, distribution of per-TCR predicted scale for TCRs whose peptide is the same as cluster’s dominant peptide ("matches dominant peptide") vs. peptide is not the dominant peptide ("off-dominant peptide"); horizontal lines indicate group means (values shown above each panel).

To benchmark TCR embedding organization by peptide similarity, we performed Leiden clustering (k=10 graph, resolution=0.8) on each model’s embedding and scored clusters against peptide labels by Adjusted Rand Index (ARI) and Normalized Mutual Information (NMI); all models used the same fold-3 split (75,884 reference, ∼23,400 test TCRs). PRISM led on both metrics and sets (Fig. 2c). On the reference set PRISM reached ARI=0.109 (95% CI 0.106–0.112) and NMI=0.287 (0.284–0.289), exceeding every baseline (Mann-Whitney U, Bonferroni p<0.05): SCEPTR (0.028, 0.129), TCR-BERT (0.004, 0.086), CDR3 Levenshtein (−0.002, 0.091), ESM2 (0.001, 0.061). On test, PRISM again led with ARI=0.043 (0.040–0.046), NMI=0.168 (0.164–0.172) versus SCEPTR (0.023, 0.151), TCR-BERT (0.005, 0.100), CDR3 Levenshtein (0.003, 0.095), and ESM2 (0.002, 0.072).

PRISM generates multiple high-purity clusters. On the reference set, a 5,918-TCR cluster was 90.2% YVLDHLIVV, a 3,406-TCR cluster 84.5% GLCTLVAML, and a 1,718-TCR cluster 76.7% SEHDYQIGGYTEKW; smaller clusters reached comparable purity (e.g., 100% RPIIRPATL, n=15). These results held up on the test set, YVLDHLIVV stayed the purest large cluster (67.7%, n=1,830), followed by SEHDYQIGGYTEKW (58.6%, n=596). Clusters dominated by KLGGALQAK and HTTDPSFLGRY were the exception (10–24% purity); both recur as secondary peptides in other clusters, so their TCRs disperse through the embedding rather than forming one population. Several mixed clusters reflected real epitope similarity: one merged CTPYDINQM, TTPESANL, and STPESANL (Lentivirus peptides, the latter two differing by one residue), and another split evenly between FRDYVDRFYKTLRAEQASQE and its substring RFYKTLRAEQASQ. In cases where epitope sequences overlap, PRISM embeds their TCRs together, forming a single shared cluster rather than separating by antigen.

Cluster purity also tracked PRISM’s learned scale parameter that reflects uncertainty. Alongside each embedding, the per-receptor scale parameter reflects how well an individual TCR is resolved within the embedding, high where a receptor sits in a tight, coherent neighborhood and low where it does not. Mean predicted scale per leiden cluster correlated with cluster purity (reference r = 0.82, test r = 0.80), and within clusters, TCRs matching the dominant peptide carried higher scale than those specific to other peptides (reference 23.7 vs 11.9; test 21.7 vs 12.0; Fig. 2d). The learned scale thus flags well-resolved regions of the embedding without reference to peptide labels.

### PRISM matches or exceeds existing methods on held-out and independent benchmarks

Prediction performance was assessed using area under the receiver operating characteristic curve (AUROC), and partial AUROC at a false-positive rate of 0.1 (pAUROC@0.1), each computed per peptide and averaged (macro AUROC, macro pAUROC@0.1) to account for dataset imbalance. Because standard 1:5 negative sampling uses a fixed set of negative peptides per TCR, which may bias the evaluation toward or against particular methods depending on the chosen negatives, we assessed robustness to different sets of negatives by precomputing scores for all TCR–peptide pairs across all models and resampling 1,000 independent 1:5 negative sets per positive TCR. On a held-out test set spanning 923 peptides, PRISM with synthetic augmentation achieved the strongest weighted macro AUROC of 0.813 ± 0.000137 (95% CI across 1,000 resamplings), followed by PRISM without augmentation (0.787 ± 0.000152), SCEPTR (0.738 ± 000083), ERGO-II (0.636 ± 000651), tcrdist3 (0.568 ± 0.000108), and NetTCR-2.0 (0.495 ± 000757) (Fig. 3a). The negligible confidence intervals confirm that the ranking is robust to the choice of negative peptides.

**Figure 3.**
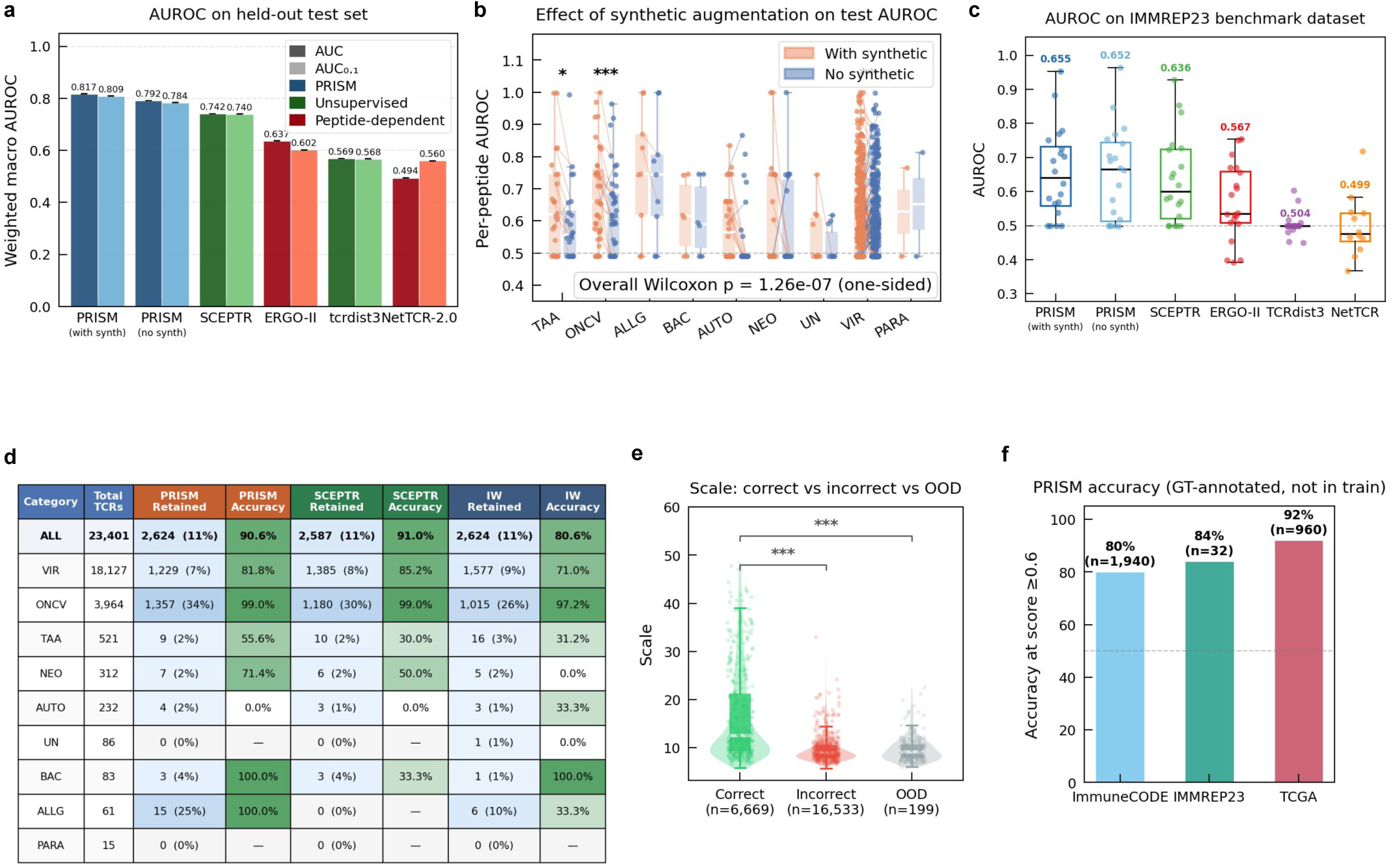
PRISM achieves accurate TCR–antigen annotation with discriminative uncertainty across benchmarks. **(a) Weighted macro AUROC on the held-out test set (923 peptides) with 95% confidence intervals from 1,000 independent negative resamplings.** PRISM with synthetic augmentation (with_synth) is benchmarked against PRISM without augmentation (no_synth) in blue, TCR sequence only models SCEPTR, tcrdist3 (green), and TCR-peptide models NetTCR-2.0 and ERGO-II (dark red). **(b) Per-peptide AUROC by antigen category for PRISM trained with versus without synthetic augmentation (orange, with synthetic; blue, no synthetic).** Categories: tumour-associated antigens (TAA), oncoviral (ONCV), allergen (ALLG), bacterial (BAC), autoimmune (AUTO), neoantigen (NEO), unknown (UN), viral (VIR), parasitic (PARA). Asterisks denote per-category significance (*P < 0.05, **P < 0.01, ***P < 0.001; one-sided Wilcoxon signed-rank); overall Wilcoxon P = 1.26 × 10⁻⁷. **(c) Macro AUROC on the independent IMMREP23 benchmark (20 peptides).** Boxplots showing per peptide AUROC across methods; overlaid text is the macro AUROC value. Dashed line, random classifier (0.50). **(d) Annotation accuracy on held-out test set at matched 11% retention for PRISM, SCEPTR, and ImmuneWatch DETECT (IW).** For each method, "Retained" columns give the number and percentage of the category’s total TCRs included; "Accuracy" columns give top-1 annotation accuracy on the retained set. Em-dashes indicate categories with no retained receptors. PRISM shows higher accuracy on the under-represented TAA, NEO, and ALLG categories. (**e) Distribution of the learned scale parameter.** Violin plot of correctly annotated TCRs (n = 6,669), incorrect (n = 16,533) or out of distribution (OOD) (n = 199) TCRs (***P < 0.001, Kruskal–Wallis test with pairwise comparisons). White lines indicate medians; box-and-whisker overlays show interquartile range and 1.5 × IQR. **(f) PRISM annotation accuracy on three orthogonal datasets at a fixed kNN confidence threshold of 0.6, restricted to receptors with ground-truth annotations not present in training:** ImmuneCODE (MIRA-assay SARS-CoV-2 TCRs; n = 1,940), IMMREP23 (n = 32), and TCGA (n = 960). Dashed line, 50%.

On the independent IMMREP23^26^ benchmark of 20 epitopes, PRISM again attained the highest overall macro AUROC and macro area under the precision-recall curve (AUPRC) (0.655; macro AUPRC 0.417) (Fig. 3b), exceeding PRISM without augmentation (AUROC 0.652, AUPRC 0.416), SCEPTR (0.636, 0.398), ERGO-II (0.567, 0.260), NetTCR-2.0 (0.499, 0.212) and tcrdist3 (0.504, 0.177), and substantially exceeded the no-skill baseline defined in Methods (AUPRC 0.17). Performance across both held-out and independent benchmarks indicates that PRISM generalizes to TCRs outside of training data.

### Structure-guided data augmentation improves representation learning for underrepresented epitopes

The reference is dominated by common viral and oncoviral epitopes, many represented by thousands of receptors, while tumor-associated, neoantigen and other classes are sparsely sampled (Supplementary Fig. 1a,c). PRISM addresses this imbalance with `airr-gen`, which generates synthetic receptors for underrepresented antigens by mutating CDR non-interface positions identified from corresponding predicted TCR–pMHC structures and retains variants whose predicted binding free energy and fold stability remain within defined tolerances (Methods). Synthetic receptors are sampled to constitute the majority of the training data for the sparsest classes and a minority for the well-sampled ones (Supplementary Fig. 2). Across all peptide specificities, the average performance gain associated with augmentation was modest, reflecting the strong representation of many common antigens within existing reference datasets. Improvements were concentrated among epitopes with sparse experimental data : per peptide AUROC improved significantly within the tumor associated antigens (TAA) (+5.6%, p = 2.6 × 10⁻²), oncoviral (ONCV) (+5.5%, p = 3.5 × 10⁻⁴) and viral (VIR) (+1.8%, p = 5.7 × 10⁻³) categories, with an overall one-sided Wilcoxon p value of 1.26 × 10⁻⁷ across evaluated peptides (Fig. 3c). Structure guided data augmentation therefore improves annotation precisely where experimentally characterized receptors are scarcest.

### An intrinsic per-receptor uncertainty distinguishes well supported from out of distribution receptors and supports accurate predictions

PRISM associates each annotation with two measures of reliability; 1) the kNN confidence score based on neighborhood agreement and 2) the learned scale parameter reflecting embedding uncertainty. The first derives from the retrieval itself: At inference, antigen assignments are generated by distance-weighted k-nearest-neighbor (kNN, k = 10) querying the antigen-labelled reference atlas across a 5-fold ensemble. Retaining receptors above increasing neighbourhood-agreement thresholds raised top-1 accuracy monotonically, from 28.5% over all predictions to 84% over the most confident decile, establishing that assignments backed by consistent local evidence are substantially more likely to be correct.

At a test–set retention proportion matched across methods (11%, PRISM kNN confidence >= 0.6), PRISM achieved overall accuracy comparable to SCEPTR overall while exceeding it on the underrepresented tumor-associated (55.6% vs 30%), neoantigen (71.4% vs 50%) and allergen (100% vs –, no TCRs retained) categories, and exceeded the ImmuneWatch DETECT^27^ platform overall (90.6% vs 80.6%) as well as on those categories (Fig. 3d). High accuracy using PRISM kNN confidence threshold of 0.6 threshold was observed on TCRs that had a peptide label but were outside the training data across ImmuneCODE^28^ (MIRA-assay based SARS-CoV-2 TCR, accuracy - 80%(, IMMREP23 benchmark (accuracy - 84%) and TCGA (The Cancer Genome Atlas bulk RNA-seq inferred TCRs, accuracy = 92%) showing that confidence-gated annotation generalizes across datasets (Fig. 3f).

The second reliability measure derives from the embedding itself: a per receptor scale parameter *s*, learned jointly with the representation. Scale was higher for receptors in tight, coherent clusters and was uncorrelated with class size in the training embedding, so it reflects local embedding structure rather than how many training examples a peptide has (Supplementary Fig. 3a,b). In the held-out test set, scale distinguishes correct, incorrect, and out-of-distribution predictions, reflecting kNN confidence but also distinguishes receptors associated with antigens that are absent from the training data. Scale was highest for correctly annotated TCRs (n = 6,669, mean = 16.47 ± 9.50), lower for incorrectly annotated TCRs (n = 16,533, mean = 9.77 ± 3.22), and lowest for receptors whose antigens were absent from training (n = 199, mean = 9.66 ± 2.73; Kruskal–Wallis P < 0.001, all pairwise comparisons (Fig. 3e).

### PRISM attention identifies CDR3 residues forming hydrophobic contacts with the peptide that shape the learned embedding

Per-position attention weights from each PRISM encoder branch provide a direct readout of the residues used to construct the embedding. Within each branch, the self-attention layer reweights positional contributions before the convolution stack and global pooling, so attention reflects how the encoder weighs the CDR region rather than how it scores any particular peptide. Averaged across the 398 training peptides with sufficient receptor support (minimum of 50 training TCRs), attention was distributed unevenly between the branches (Fig. 4a).

**Figure 4.**
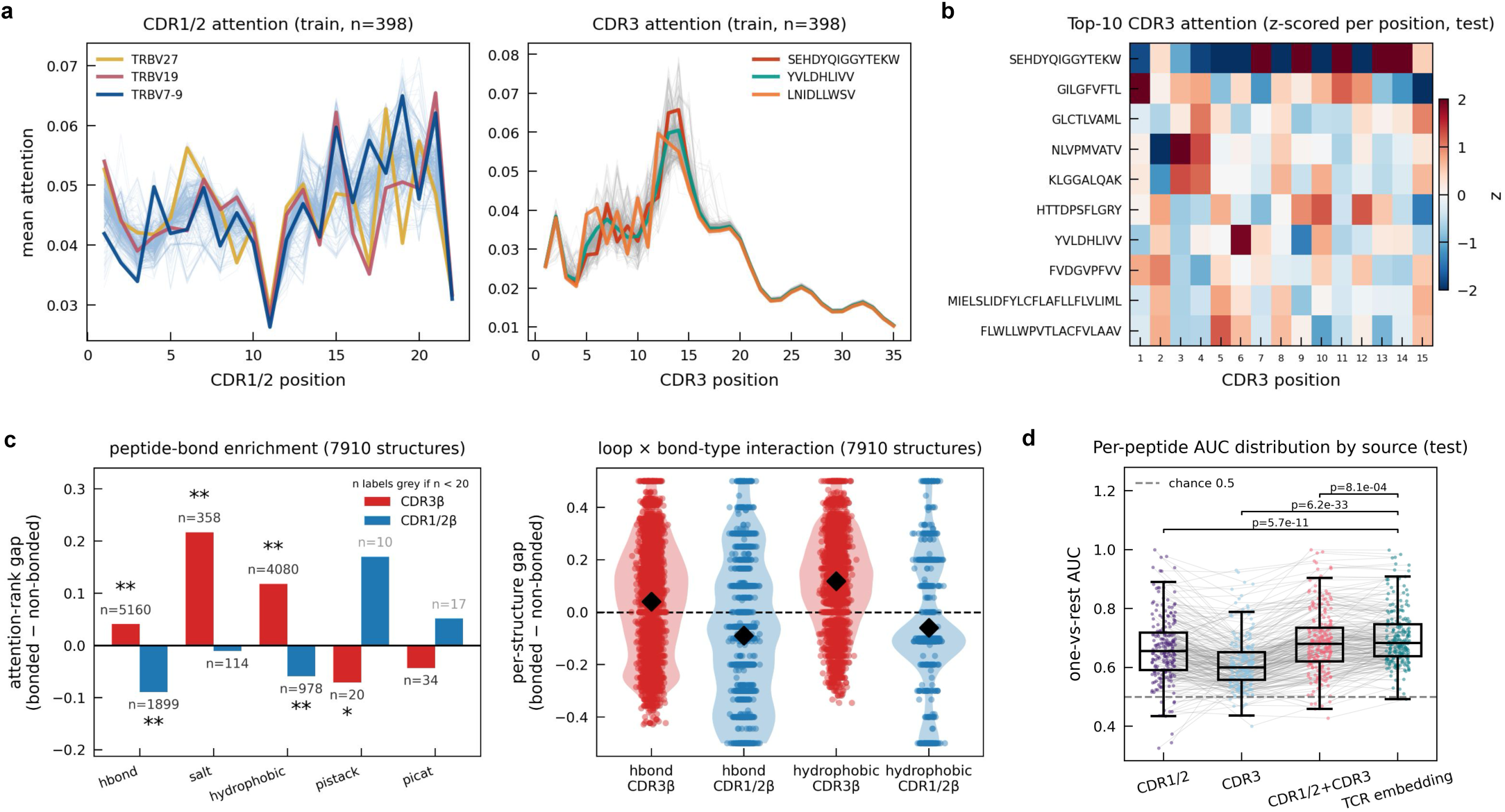
Attention is concentrated in CDR3, tracks peptide-bond residues and informs embedding that carries highest specificity signal. **(a) Mean per-position attention in CDR1/2 (left, blue) and CDR3 (right, red) for the training set (n = 398 peptides; one trace per peptide).** CDR1/2 attention is comparatively flat across positions, whereas CDR3 attention peaks in the central-to-C-terminal loop region (positions ∼12–15). **(b) Per-position CDR3 attention for the ten most frequently observed peptides in the training set** (subsampled to 500 TCRs per peptide), z-scored within each CDR3 position to highlight peptide-specific deviation from the position-wise mean. Rows, peptide epitopes; columns, CDR3 position (1–15); color, z-score (red, above-average attention; blue, below-average). **(c) Attention enrichment on peptide-bonding residues, computed from predicted TCR–pMHC structures.** Bonds are specific TCR–peptide interactions identified by PLIP (hydrogen bond, salt bridge, hydrophobic, π-stacking, π-cation), independent of spatial proximity. Within each structure, loop residues were ranked by attention weight (scaled to [0,1]); the attention-rank gap is the mean rank of residues forming a given bond to the peptide minus the mean rank of residues not forming that bond (positive, attention favors bond-forming residues). Left, mean gap per bond type for CDR3β (red) and CDR1/2β (blue); n, structures containing ≥ 1 bond of that type (gray where n < 20). Right, per-structure gap distributions for the two best-powered bond types (hydrogen bond, hydrophobic), split by loop, where points are structures and diamonds are means. CDR3β attention is enriched on salt bridges and hydrophobic bonds, and weakly on hydrogen bonds, whereas CDR1/2β gaps are negative, reversing the sign of the enrichment between loops; salt bridges are omitted from the right panel (too few for a per-structure distribution). Wilcoxon signed-rank test against zero; ***P < 0.001, *P < 0.05. **(d) Specificity classification performance on the held-out test set from multinomial logistic regression trained on the training set, restricted to the 398 peptides with ≥ 50 training TCRs.** Box plot of per-peptide one-vs-rest AUC by input source (CDR1/2, CDR3, CDR1/2+CDR3, TCR embedding); boxes show median and interquartile range, grey lines connect the same peptide across sources, and the dashed line marks chance (0.5). P-values are from Bonferroni corrected Wilcoxon signed rank test.

CDR1β/CDR2β attention was comparatively flat across positions, whereas CDR3β attention was concentrated in the central-to-C-terminal segment of the loop (approximately positions 12–15). This was not a fixed positional prior: z scoring CDR3β attention within each position to remove the population-average profile, revealed peptide-specific deviations for most frequent test peptides (Fig. 4b), indicating that the encoder shifts its positional focus with antigen context rather than attending to identical residues for every receptor.

To test whether this attention tracks the chemical basis of recognition, we profiled TCR–peptide interactions with PLIP^29^ across 162 experimentally determined and 7910 predicted TCR–pMHC structures and compared attention on the CDR residues forming each interaction type against the remaining loop residues (Fig. 4c). In predicted structures, CDR3β attention was most strongly enriched on salt bridges (mean attention-rank gap +0.22) and hydrophobic contacts (+0.12); enrichment on hydrogen bonds was statistically significant but negligible in magnitude (+0.04). This enrichment was specific to bond type: among CDR3β-core residues, attention correlated most strongly with hydrophobic-contact count (Spearman ρ = +0.21) and was uncorrelated with either distance to the peptide or untyped peptide contact (Supplementary Fig. 4a,b). CDR1/2β gaps were negative (hydrogen bond −0.09, hydrophobic −0.06), so the germline loops attend away from the peptide-contacting residues, reversing the sign of the enrichment between loops (Fig. 4c; Supplementary Fig. 4c); the per-structure distributions confirmed that the CDR3β hydrophobic enrichment is a shift of the whole population rather than a few structures (Fig. 4c, right). Enrichment of CDR3β attention on salt bridges and hydrophobic contacts is consistent with the structural basis of recognition: salt bridges provide directional, charge-specific readout of the peptide^30,31^, whereas hydrophobic packing forms the bulk of the buried interface and is its principal energetic determinant^32,33^. The same enrichment of CDR3β attention on salt bridges (gap = +0.13, P = 0.009) and hydrophobic contacts (+0.06, P = 0.002) was independently replicated in 162 experimentally determined crystal structures from TCR3d^34^ (Supplementary Fig. 4d,e), confirming the signal reflects genuine TCR–peptide interface bond chemistry, and not an artifact of predicted structures. Attention thus concentrates on the CDR3 residues that underlie both the specificity and the stability of the TCR–peptide interface.

To test how much each component of the representation contributes to discriminating peptides, we used each branch’s attention, pooled representation and final embedding as inputs to multinomial logistic regression over the same 398 peptides. All four sources classified peptides above chance (Fig. 4d). The CDR1/2β representation alone (mean per-peptide AUROC 0.663 ± 0.114) was a stronger standalone classifier than CDR3β alone (0.614 ± 0.089); concatenating the two improved performance (0.687 ± 0.108), and the full embedding was strongest (0.703 ± 0.095). Germline context out-predicts CDR3β reflects peptide-associated V-gene usage in the training set, which the CDR1/2β branch reads directly; CDR3β adds complementary, antigen-discriminating signal (+0.024 over CDR1/2β alone), and the two branches together recover nearly all of the full embedding’s performance.

## Discussion

PRISM addresses a setting that recurs in disease, vaccination, and longitudinal monitoring: repertoires whose receptors carry no known specificity and for which no candidate antigens are available. Its premise is that a representation organized by antigen, rather than by sequence similarity, is required, because the mapping between sequence and specificity is not monotonic—near-identical receptors can recognize distinct antigens, and receptors converging on the same antigen can share little sequence. PRISM learns such a representation by supervised metric learning, embedding TCRβ sequences so that shared specificity, not sequence proximity, defines neighborhoods. This supports two complementary operations: a receptor can be annotated by retrieving its nearest labeled neighbors, together with an assessment of whether that assignment is well supported; and the neighborhoods themselves can be interrogated, recovering groups of sequence-diverse receptors that similarity-based methods may leave dispersed. On a 923-peptide held-out set and the independent IMMREP23 benchmark, annotation matches or exceeds sequence-based models, and receptors outside the reference are abstained on rather than forced into known classes.

Two properties distinguish PRISM as a representation for repertoire-scale annotation. The first is that annotation proceeds by retrieval against a labeled reference rather than by classification into a fixed antigen set: a receptor is placed only on the evidence of its nearest neighbors, so receptors without well-matched neighbors are left unannotated rather than assigned to the closest available class. The per-receptor uncertainty that PRISM learns alongside the embedding makes this principle operational—it reflects whether a receptor occupies a well-supported region of antigen space, separates correct from incorrect assignments, and is lowest for receptors whose antigens are absent from the reference. Each annotation therefore carries an estimate of its own reliability, and the receptors the model cannot place are identified as readily as those it can. This is what renders annotation tractable at the scale of a repertoire.

The second property concerns the structure of the training data. Public reference databases are dominated by immunodominant viral epitopes presented on common HLA alleles, whereas the tumor-associated and neoantigen specificities of greatest translational interest are sparsely represented or absent—an imbalance under which a model attains competence only over a privileged region of antigen space. PRISM counters this with structure-guided synthetic receptors that diversify non-interface positions while holding fixed the contacts that define recognition. The resulting gains concentrate in the underrepresented categories rather than distributing evenly across antigens already well covered, so augmentation does not simply raise mean performance; it relocates where the model is competent, extending reliable annotation into regions of antigen space that experimental data leave thin.

Whether an antigen-organized embedding captures genuine determinants of recognition, rather than sequence covariates, is an interpretability question that PRISM’s attention weights allow us to address directly. Across predicted and experimentally determined TCR–pMHC interfaces, CDR3β attention localizes to the residues forming salt bridges and hydrophobic contacts with the peptide—the charge-complementary contacts that confer sequence-specific readout and the apolar packing that constitutes the bulk of the buried interface and its principal energetic determinant—while the germline-encoded CDR1/2β loops are directed away from the peptide. This localization is governed by the chemistry of the contact rather than by spatial proximity: attention is no greater on residues near the peptide than on distal ones, and it discriminates among interface residues by the bond they form, not by their distance. The embedding is therefore organized around the chemical determinants of peptide recognition rather than sequence features that need not track specificity. For a model whose assignments are hypotheses, this is the relevant assurance: PRISM’s neighborhoods reflect how receptors engage the peptide, not how closely their sequences resemble one another.

PRISM’s annotations remain bounded by its reference, which inherits the biases of public TCR–antigen data; confidence-gating attenuates this dependence but does not abolish it, and even high-confidence assignments are candidates awaiting experimental validation. PRISM was deliberately constructed as a β-chain model, because paired-chain and HLA-resolved data remain scarce and most repertoire sequencing reports the TCRβ chain alone. These constraints are open problems for the field, and an antigen-organized, confidence-aware representation is a natural scaffold into which paired α-chain, peptide–MHC structural, and HLA-restriction information can be incorporated as it accrues.

The central contribution of PRISM is the antigen-organized representation itself, which serves annotation and discovery alike. By grouping receptors according to shared specificity, PRISM can surface receptor sets that recur across disease repertoires—candidates for shared, potentially disease-associated specificities, against antigens that no reference yet contains. The attention weights offer an interpretable view into the organization of receptors and can be leveraged when examining possible antigen specific receptor groups. We anticipate that such a representation will become a substrate for discovery across immune monitoring, vaccine-response profiling, and the identification of disease-associated T cells, transforming repertoire sequencing from a catalogue of receptors into a map of what they recognize.

## Methods

### Dataset curation and preprocessing

TCR–antigen binding data were obtained by aggregating and harmonizing TCR–pMHC interaction records from public sources – IEDB, VDJdb, McPAS-TCR, TCR3d, CEDAR (all last accessed December 2025) and patents from PATCRdb (230,189 total records; 135,126 unique CDR3β sequences; 2,602 unique epitopes). Along with sequence data, PDB files for x-ray crystallography derived structures from TCR3d were downloaded for analysis (184 structures with human TCR beta chain). Quality control filtering retained 204,206 records (133,218 unique Vβ–CDR3β clonotypes across 2,009 peptides) by excluding entries missing CDR3β sequence, V gene assignment, or peptide annotation, and retaining only TCRs encodable by the CDR1β–CDR2β–CDR3β concatenation scheme. After deduplication on (Vβ–CDR3β, peptide) pairs, 4,663 cross-reactive TCRs (those binding >1 peptide) were removed. Only peptides with at least 2 unique TCR sequences were retained, yielding 116,303 unique TCR–peptide pairs across 1,159 peptides. Each TCR β-chain was encoded by concatenating the amino acid sequences of its CDR1β, CDR2β, and CDR3β regions, represented as integer indices over a 22-symbol vocabulary (20 standard amino acids, a padding token “X”, and a gap character “-” to a fixed-length input vector of 57 positions. To quantify peptide-level sequence similarity for use in loss weighting, pairwise BLOSUM62 substitution scores were computed between all peptide pairs and normalized to the range [0, 1] to produce a pre-computed peptide similarity matrix. The dataset was partitioned using a hybrid peptide- and TCR-level split strategy. First, 10 peptides (10–50 TCRs each) were held out as an out-of-distribution (OOD) test set (198 TCRs). The remaining 1,149 peptides were split at the TCR level using stratified 80/20 sampling, producing a training set (92,884 TCRs) and a seen-peptide test set (23,221 TCRs) with non-overlapping TCR sequences. The combined test set (23,419 TCRs across 923 peptides) includes both seen-peptide and OOD partitions. The training set was further divided into 5-fold cross-validation splits (90/10 stratified by peptide, ∼75,883/17,000 TCRs per fold); only peptides with ≥ 50 TCRs contributed to validation folds, while lower-frequency peptides were always included in training. For AUROC evaluation, negative pairs were generated by randomly assigning peptide labels other than the true peptide to TCR sequences at a 1:5 positive-to-negative ratio, like done in IMMREP23^26^.

### Structure-guided synthetic data augmentation

To address the class imbalance inherent in curated TCR-epitope databases, where a small number of well-studied viral epitopes dominate the training distribution, PRISM incorporates a structure-guided synthetic data augmentation pipeline, `airr-gen`. For 28,086 TCR–pMHC complexes in the training set where TCR alpha chain, TCR beta chain, peptide, and MHC allele annotations were available, TCR-pMHC complex structures were predicted using TCRmodel2^35^. Each top-ranked (rank-0) model was scored and marked PASS when all of the following held: TCR–pMHC interface confidence (ipTM) ≥ 0.8, AlphaFold assembly confidence (ipTM) ≥ 0.8, global complex confidence (pTM) ≥ 0.6, and predicted backbone accuracy (pLDDT) > 80. Of 28,086 structures, 10,428 passed quality filtering. For each passing structure, interface contacting residues, identified as TCR CDR positions with any atom within 5 Å of any peptide, MHC, or paired TCR-chain atom, together with conserved CDR3 residues were locked. Synthetic variants were generated by introducing single and double amino-acid substitutions, under a neutral substitution strategy, at the remaining non-interface CDRβ positions.

Candidate variants were evaluated with FoldX 5.1^36^ under its default force-field parameters with energies reported in kcal/mol. The wild-type complex and a separately extracted free TCR (α and β chains only) were each minimized once with RepairPDB, and these repaired structures defined the reference state for all energy differences. Mutant structures were built from the repaired templates with BuildModel using a single model per variant (numberOfRuns = 1): full-complex mutants from the repaired complex and free-TCR mutants from the repaired TCR. Folding free energy (G) was computed for every structure with the Stability command, and the TCR–pMHC interaction energy was computed for the full complexes with AnalyseComplex, partitioning the assembly into a TCR group (α, β) and a pMHC group (peptide, MHC). For each variant, the change in complex folding stability, the change in free-TCR folding stability, and the change in interaction energy were each taken as the mutant-minus-wild-type difference of the corresponding FoldX energy. Variants were retained only when |ΔΔG_stability(complex)| < 1 kcal/mol, |ΔΔG_stability(TCR)| < 1 kcal/mol, and |ΔΔG_interaction| < 0.5 kcal/mol (Equation 5), ensuring that each synthetic TCR preserved both the TCR fold and the binding interface (Supplementary Fig. 2b). The CDR1β, CDR2β, and CDR3β sequences of passing variants were extracted and encoded identically to observed TCRs. For peptides with more than 100 observed TCRs, synthetic TCRs were capped at 150 per peptide; for peptides with ≤ 100 observed TCRs, all available synthetic variants were retained (Supplementary Fig. 2c). This yielded 115,216 synthetic TCRs across 306 peptides, augmenting the training data of 92,884 observed TCRs across 1,149 peptides.

### Model Architecture

PRISM encodes TCR β-chain sequences through a dual-branch neural network that processes the CDR1β/CDR2β and CDR3β regions separately before fusing them into a shared latent space. Each branch consists of a learned 22 dimensional amino acid embedding, sinusoidal positional encoding, a self-attention layer with L1-regularized query, key and value projections that learns to weight positional contributions within each CDR region, followed by three stacked one-dimensional convolutional layers with decreasing kernel sizes (7, 5, 3), batch normalization, ReLU activation, dropout (rate 0.2), and max pooling. The output of each branch is summarized by global average pooling to produce a fixed-length representation. The two branch representations are concatenated and projected through two additional convolutional layers with batch normalization and dropout, then flattened and passed through a linear projection to yield a 128-dimensional embedding *z*, L2-normalized to the unit hypersphere.

A separate intrinsic uncertainty head branches from the same fused representation: a two-layer multi-layer-perceptron (128 and 64 units, ReLU, batch normalization, dropout) followed by a sigmoid-activated output linearly scaled to [5, 50], producing a per-TCR scalar *s* that reflects confidence in embedding quality. The model additionally exposes per-position attention weights from each branch, enabling post-hoc interpretation of CDR position contributions to the embedding.

### Training objective

PRISM is trained with a combined metric-learning objective: a peptide-similarity-weighted supervised contrastive loss and a ScaleFace loss that simultaneously learns embedding geometry and per-sample confidence.

#### Supervised contrastive loss

For a batch of *B* TCRs with L2-normalized embeddings *z* and peptide labels *y*, we extend the supervised contrastive objective of Khosla et al^37^. to graded inter-class similarity:

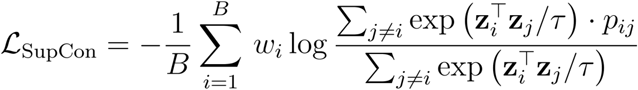

where *p_ij_* is the BLOSUM62-derived peptide similarity between the target peptides of TCRs *i* and *j* if >= 0.7 and 0 otherwise, *τ* = 0.3, and *w_i_* is a per-sample weight (*w_i_* = 1.0 for experimentally validated TCRs, *w_i_* = 1.0 for synthetic TCRs). In standard SupCon, all same-class pairs are treated as binary positives; here, every pair in the batch contributes to the numerator if its target peptides are highly similar, pulling TCRs together continuously rather than discretely. This is particularly important for TCRs targeting related epitopes (e.g. peptide variants or HLA-restricted variants of the same antigen), which should occupy nearby but not identical positions in embedding space.

#### ScaleFace loss

Following Kail et al^38^., we introduce a trainable class weight matrix **W** ∊ ℝ*^C^*^× 128^ (one normalized prototype per epitope class) and a per-TCR scale *s_i_* ∊ [5, 50] predicted by the uncertainty head. The margin-adjusted, scale-modulated logit, 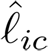 for sample *i* and class *c* is:

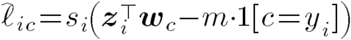

where *m* = 0.35 is the cosine margin. The ScaleFace loss is:

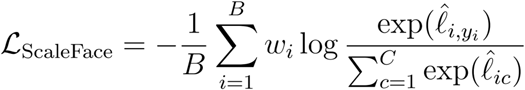

A high predicted *s_i_* sharpens the softmax, strengthening the gradient for confident predictions; a low *s_i_* flattens it, reducing the penalty for uncertain ones, coupling classification and calibration in a single forward pass. Uncertainty per TCR is computed as 1/*s_i_* ranging from 0.02 to 0.20.

Combined objective:

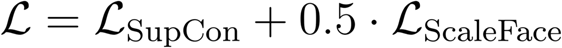

Encoder weights, the prototype matrix W and uncertainty head are optimized jointly.

The model was trained using the Adam optimizer with a learning rate of 0.001 and a batch size of 8000. Five folds of training ran for up to 1,000 epochs each with early stopping monitored on a cross fold validation split of 10% of training TCRs, stratified by peptide. Early stopping was based on k-nearest-neighbor Precision@5 (kNN P@5) computed on the validation set every epoch, i.e., the fraction of the five nearest neighbours in embedding space sharing the query TCR’s epitope label. This directly reflects embedding quality under a metric-learning objective. The checkpoint with the highest kNN P@5 was retained, and training was halted if no improvement was observed for 100 consecutive epochs. L1 regularization (λ = 1 × 10⁻⁶) was applied to all learned weights and dropout (rate 0.2) throughout the encoder and uncertainty head. The prototype matrix W was initialized from a standard normal distribution (σ = 0.1) and optimized jointly with all encoder parameters. All experiments used a fixed random seed of 42 and were run on a single NVIDIA A100 GPU.

### Inference and scoring

At inference, the query TCR is encoded to a 128-dimensional L2-normalized embedding and its k=10 nearest neighbors are retrieved from each fold’s epitope-labeled reference set by cosine distance using faiss (v1.11.0)^39^. For a query embedding *z_q_* and neighbor *z_j_*, cosine distance is *d_cos_*(**z***_q_*, **z***_j_*) = 1 - **z***_q_* **z***_j_*, and each neighbor is weighted by the reciprocal of its distance to the query,

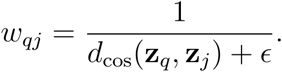

The per-fold score for peptide *p* is the distance-weighted fraction of the *k* neighbors 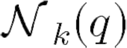 carrying that label,

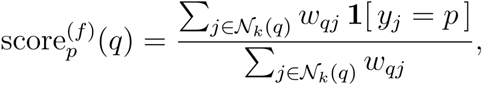

so that scores form a distribution over the labels present among the neighbors. Final scores are averaged across the five folds,

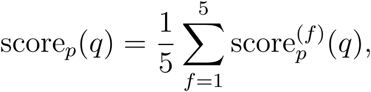

and the top-1 prediction is the highest-scoring peptide, with confidence score*_p_(q)*. A prediction is retained only when the top score exceeds a minimum threshold and the per-TCR uncertainty falls below a maximum, otherwise the receptor is abstained on. Recommended operating thresholds are derived from the score and scale distributions of held-out data (Results).

### Embedding quality evaluation with Leiden clustering

Embedding quality of PRISM was evaluated against SCEPTR, TCR-BERT, ESM2 and CDR3 Levenshtein distance. For each model and split, generated embeddings (or CDR3β sequences for the Levenshtein baseline) were used to construct a k-nearest-neighbor (k=10) similarity graph: cosine similarity via exact inner-product search (faiss v1.11.0) for embedding-based models, and pairwise Levenshtein edit distance (rapidfuzz v3.14.3) for the sequence-based baseline. Community detection was performed with the Leiden algorithm (leidenalg, RBConfigurationVertexPartition, resolution parameter = 0.8, seed = 0) on the resulting weighted graph. Cluster assignments were scored against ground-truth peptide-specificity labels using the adjusted Rand index (ARI) and normalized mutual information (NMI). To enable a matched comparison, all models’ "reference" (train) sets were restricted to the same 75,884 TCRs comprising PRISM fold3’s training split; the test set (23,401 TCRs) was shared across models without modification.

Uncertainty was estimated by bootstrap resampling: for each model and split, TCRs were resampled with replacement (n = 1,000 iterations) and ARI/NMI recomputed on the resampled cluster/peptide-label pairs without rerunning Leiden clustering, yielding an empirical sampling distribution per model. 95% confidence intervals were taken as the 2.5th–97.5th percentiles of this distribution. To test whether each baseline model differed significantly from PRISM, a two-sided Mann–Whitney U test was applied between each baseline’s bootstrap distribution and PRISM’s, separately for each split and metric (ARI, NMI).

### Benchmarking

We evaluated PRISM on benchmark using identical train/test splits used for PRISM. Models were grouped into two tracks based on their input requirements. Track 1 (TCR-only models) included β-SCEPTR (v1.2.0), and tcrdist3 (v0.3). β-SCEPTR was evaluated using its pretrained model without retraining; binding scores were derived from distance-weighted k-nearest-neighbor (k=10) classification in embedding space, where the score for a test pair (TCRᵢ, peptideⱼ) was computed as the weighted fraction of the k nearest training positives (by cosine distance) that bind peptideⱼ, with weights inversely proportional to distance. The same KNN scoring protocol was applied to tcrdist3 (using tcrdist TCR distances). Track 2 (TCR+peptide models) included ERGO-II and NetTCR-2.0. ERGO-II and NetTCR-2.0 were retrained from scratch on the training data using their published protocols to prevent data leakage from pre-trained weights; ERGO-II was trained for 50 epochs using its LSTM architecture, while NetTCR-2.0 was trained on CDR3β sequences restricted to 9-mer peptides (its maximum supported peptide length). All models were evaluated on the held-out test set with negatives generated at a 1:5 positive-to-negative ratio by random peptide reassignment, consistent with the ratios used in the original NetTCR-2.0 and ERGO-II publications. Performance was measured by AUROC and partial AUROC at a false positive rate of 0.1, computed per peptide and reported as macro averages. To obtain confidence intervals for per-peptide AUROC, and pAUROC@0.1, we resampled 1,000 independent 1:5 positive-to-negative evaluation sets by drawing 5 random negative peptides per positive TCR in each iteration. Metrics were computed per peptide and aggregated as weighted (by number of positive TCRs) macro averages, with 95% confidence intervals derived from the 2.5th and 97.5th percentiles of the resampled distributions. Additional evaluation was performed on the independent IMMREP23 benchmark (635 TCRs, 20 epitopes), using macro AUC as the primary metric.

We benchmarked PRISM’s accuracy of top-1 peptide prediction against SCEPTR and ImmuneWatch DETECT. ImmuneWatch DETECT v1.4.7 (academic-use license) was deployed locally via a docker container. The held-out test set (23,219 TCRs) and training set (92,884 TCRs) used for all methods were formatted in AIRR-compliant bulk repertoire format, including the required junction_aa, v_call, and j_call fields. To enable a fair comparison with SCEPTR and PRISM, the training repertoire was supplied as the reference database using the -d option, allowing DETECT to annotate the held-out TCRs against the same set of known antigen-specific receptors available to the other methods.

All evaluations were conducted using TensorFlow 2.21.0, PyTorch 2.12.0, and scikit-learn 1.6.1.

### External validation datasets

Three independent datasets were used to evaluate PRISM’s prediction accuracy on TCRs not present in the training set. First, ImmuneCODE; TCR β-chain sequences with experimentally validated SARS-CoV-2 peptide specificities were obtained from the Adaptive Biotechnologies immuneACCESS platform (https://immuneaccess.com). Each TCR is annotated with one or more peptide targets determined by the Multiplex Identification of T-cell Receptor Antigen Specificity (MIRA) assay. Second, IMMREP23; The IMMREP 2023 benchmark dataset was obtained from the IMMREP23 GitHub repository, containing TCR–peptide pairs across multiple viral epitopes curated for standardized benchmarking. Third, TCGA; TCR β-chain repertoires were reconstructed from bulk RNA-seq data of The Cancer Genome Atlas (TCGA) using TRUST4, with assembled CDR3β sequences and inferred TRBV gene assignments. For all datasets, PRISM predictions requires a TRBV gene assignment (IMGT nomenclature with allele suffix, e.g., TRBV7-2*01) and CDR3β amino acid sequence as input. TRBV gene names were harmonized to IMGT nomenclature where necessary, and CDR3β sequences were filtered to retain only those beginning with cysteine (C), ending with phenylalanine (F), and containing standard amino acids. TCRs with CDR3β length exceeding 35 residues were excluded. For each TCR, PRISM’s 5-fold ensemble generated a distance-weighted k-nearest-neighbor (k=10) top-1 peptide prediction and confidence score. To evaluate accuracy, TCRs present in the training set were excluded, and only TCRs whose ground-truth peptide annotation shared at least one 5-amino-acid subsequence (5-mer) with a training peptide were retained, ensuring that predictions were evaluated against peptides within PRISM’s learned space. A prediction was considered correct if the predicted peptide shared at least one 5-mer with any of the TCR’s ground-truth peptide annotations to account for ImmuneCODE annotations being lists of overlapping peptides.

### Attention analysis

To assess the information encoded in PRISM’s learned attention weights, we extracted per-position attention scores from the CDR1/2 branch (22 positions) and CDR3 branch (35 positions) for all TCRs in the training and test set. We evaluated the predictive capacity of these attention profiles by training multinomial logistic regression classifiers (L2-regularized, 5-fold cross-validated) to predict peptide identity or V-gene identity from four feature sources: CDR1/2 attention alone, CDR3 attention alone, CDR1/2+CDR3 attention combined, and the full 128-dimensional TCR embedding. Top-1 accuracy and per-peptide one-vs-rest AUROC were computed on the held-out test data. Statistical comparisons between per-peptide AUROC distributions were performed using two-sided paired Wilcoxon signed-rank tests with Bonferroni correction for multiple comparisons. To visualize peptide-specific attention patterns, we computed mean CDR3 attention profiles per peptide (averaged across all TCRs binding that peptide, for peptides with ≥50 TCRs), z-scored per peptide across positions, and displayed as a heatmap ordered by hierarchical clustering of peptides. Per-peptide CDR1/2 and CDR3 attention line plots were generated to illustrate the diversity of positional attention profiles across peptides.

### Attention–structure concordance

Non-covalent TCR–peptide interactions were identified using PLIP^29^. Prior to detection, each high quality TCR-pMHC predicted and repaired structure as well as each crystal structure from TCR3d^34^ was protonated within PLIP using OpenBabel^40^.

Each detected interaction was assigned to its TCRβ CDR-loop residue. For each residue, three counts were tabulated: total peptide interactions; specific polar interactions (hydrogen bonds, salt bridges, and π-cation interactions); and packing interactions (hydrophobic contacts and π-stacking). Heavy-atom minimum distances from each CDR-loop residue to the peptide chain were computed from atomic coordinates. A residue was classified as peptide-proximal if it formed at least one PLIP interaction with the peptide or its heavy-atom minimum distance to the peptide was below 4.5 Å. The CDR3β core was defined as the loop excluding the four N-terminal (V-gene–proximal) and four C-terminal (J-gene–proximal) positions, removing the conserved germline framework anchors.

For each structure and loop, residues were ranked by attention weight and the ranks rescaled to the unit interval. For each feature, the Spearman correlation between its per-residue value and CDR3β attention rank was computed over the trimmed core and the full loop: packing-interaction count (hydrophobic bonds), total peptide-interaction count (any peptide bond), specific polar-interaction count (H-bonds/salt), normalized loop position (increasing toward the J-proximal C-terminus), and heavy-atom minimum distance to the peptide (spatial proximity). The attention-rank gap was defined as the mean scaled rank of one group of residues minus the mean scaled rank of the others in the same loop, computed two ways: per interaction type, residues forming a given peptide interaction vs. those not forming it, and by proximity, peptide-proximal vs. distal residues. Attention-rank gaps were computed separately for CDR3β and for CDR1/2β and, for the per-loop comparison, for CDR1, CDR2 and CDR3 individually. Significance was assessed by two-sided Wilcoxon signed-rank tests against zero; effect sizes are reported alongside p values. Interaction annotation was restricted to the TCR–peptide interface.

## Data Availability

Experimentally validated TCR–pMHC binding data were downloaded from publicly available databases: TCR3d (https://tcr3d.ibbr.umd.edu), VDJdb (https://vdjdb.cdr3.net), IEDB (https://www.iedb.org), CEDAR (https://cedar.iedb.org), McPAS-TCR (http://friedmanlab.weizmann.ac.il/McPAS-TCR/), and PATCRdb (https://patcrdb.org). The IMMREP23 benchmark dataset is available on Kaggle and on github (https://github.com/justin-barton/IMMREP23). ImmuneCODE TCRs were downloaded from immuneACCESS (DOI: 10.21417/ADPT2020COVID). TRUST4 inferred TCRs from TCGA bulk RNA-Seq were provided by Dr. Li Song.

## Code Availability

PRISM is available for non-commercial academic use only. An inference pipeline and trained model weights will be released upon publication under an academic-use license at https://github.com/jonssonlab/prism. Use for commercial purposes requires a separate license from UC Santa Cruz. Code is available to reviewers during peer review.

## Acknowledgments

We thank the UC Santa Cruz Genomics Institute for computing resources. This work was supported by a UC Cancer Research Coordinating Committee grant and the Hellman Fellows grant to V.D.J. We thank Li Song for providing the TCGA TCR data and Joshua Stuart for valuable feedback on the manuscript.

## Author contributions

DV, LM and VDJ conceived the study. DV developed the methodology, wrote the software and conducted formal analysis. DV and VDJ wrote the manuscript. LM contributed to conceptual development, model evaluation on real-world repertoire data. NR curated data for the study. AR performed PLIP analysis. VDJ supervised the work.

---

**Supplemental Figure 1.**
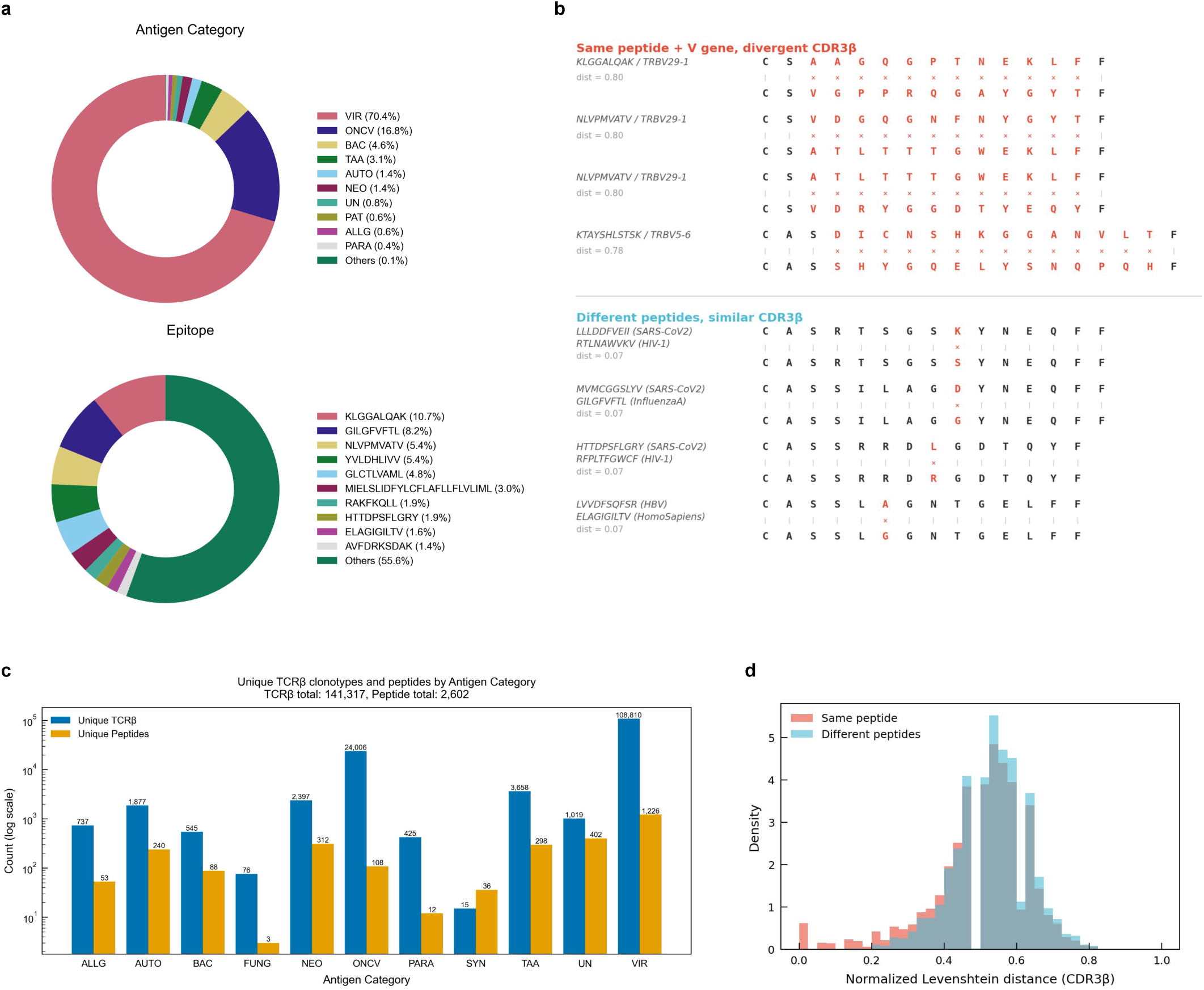
Viral peptide dominance and V-gene bias in curated experimental data. **(a) Data composition by antigen category (top) and epitope (bottom);** viral antigens dominate (VIR, 70.4%) and a few epitopes account for a disproportionate share of records (e.g., KLGGALQAK, 10.7%)**. (b) Representative TCRβ examples showing that receptors with the same cognate peptide can have divergent CDR3β (top), whereas receptors with similar CDR3β can recognize different peptides (bottom);** normalized pairwise edit distance (dist) at left, ‘|’ denote match and ‘x’ denotes mismatch. **(c) Unique TCRβ clonotypes (blue) and peptides (orange) per antigen category** (log scale; 141,317 TCRβ, 2,602 peptides total) **(d) Normalized CDR3β Levenshtein distances for sequence pairs sharing the same peptide (red) versus different peptides (blue);** the substantial overlap showing sequence distance alone poorly separates specificity.

**Supplemental Figure 2.**
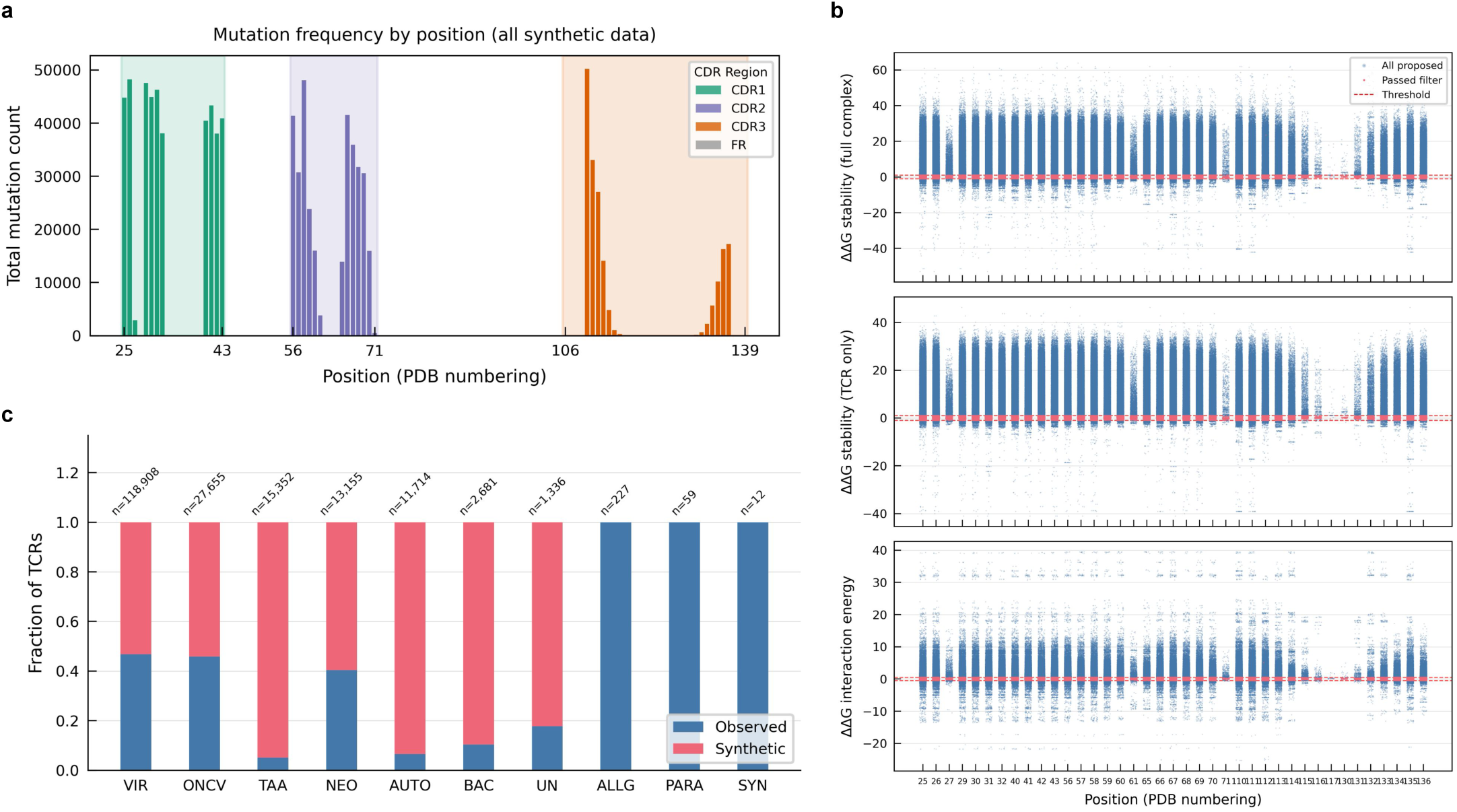
airr-gen generates structurally neutral synthetic TCRβ sequences. **(a) Positional distribution of mutations across all synthetic TCRs,** plotted by PDB residue number and shaded by CDR region (CDR1 green, CDR2 purple, CDR3 orange, framework grey). Mutations are within the CDR loops and depleted at interface-critical positions. **(b) Per-position structural quality of proposed mutations, s**cored as change in free energy (ΔΔG) relative to the parent complex for full-complex stability (top), TCR-only stability (middle), and TCR–pMHC interaction energy (bottom). Each point is a proposed mutation; the red dashed line marks the acceptance threshold and points passing the structural filter are retained. Accepted mutations cluster near ΔΔG ≈ 0, confirming that synthetic sequences are predicted to be structurally and energetically neutral**. (c) Fraction of observed (blue) versus synthetic (red) TCRs per antigen category after augmentation (n above each bar).** Augmentation is concentrated in the sparse categories — tumor-associated antigen (TAA), neoantigen (NEO), self antigen (AUTO), bacterial (BAC) and unclassified (UN), where synthetic sequences make up the majority — while well-represented viral (VIR) and oncoviral (ONCV) categories receive proportionally fewer, and categories with too few seed structures (ALLG, PARA, SYN) receive none.

**Supplemental Figure 3.**
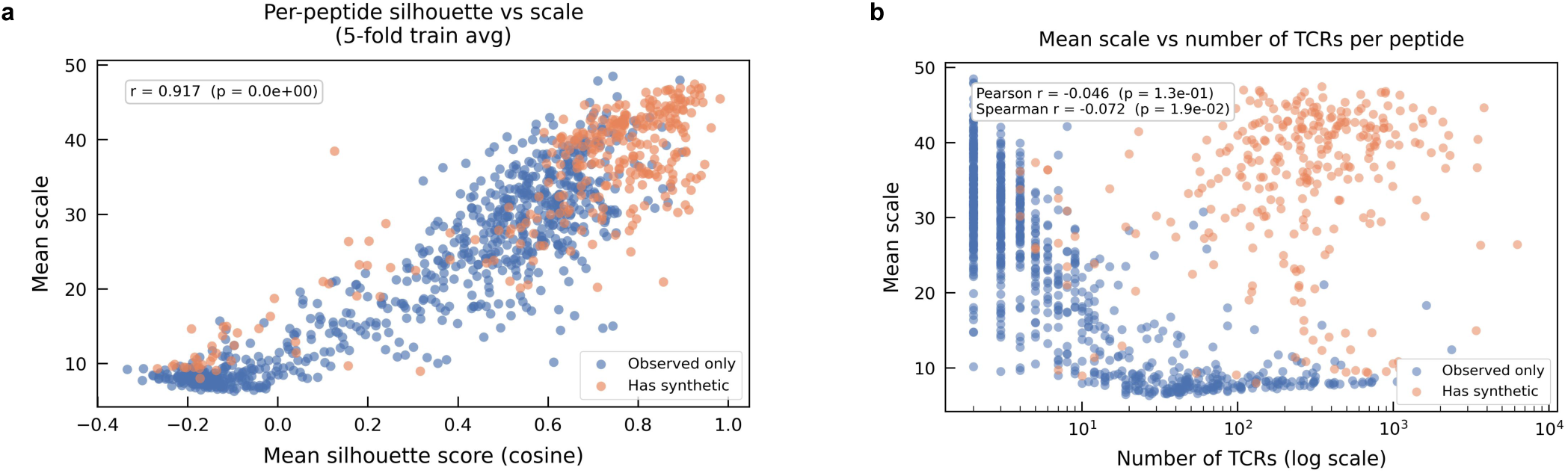
PRISM learned scale correlates with embedding separation not with class size. **(a) Mean scale per peptide vs mean silhouette score.** Scatter plot of mean silhouette score per peptide vs mean scale per peptide in the training embedding averaged across 5-folds. Spearman correlation r = 0.917. **(b) Mean scale vs number of training TCRs per peptide.** Scatterplot of number of TCRs representing a peptide in the training data vs mean scale per peptide. Spearman r = −0.046 (p = 0.13) and Pearson r = −0.072 (p = 0.012).

**Supplemental Figure 4.**
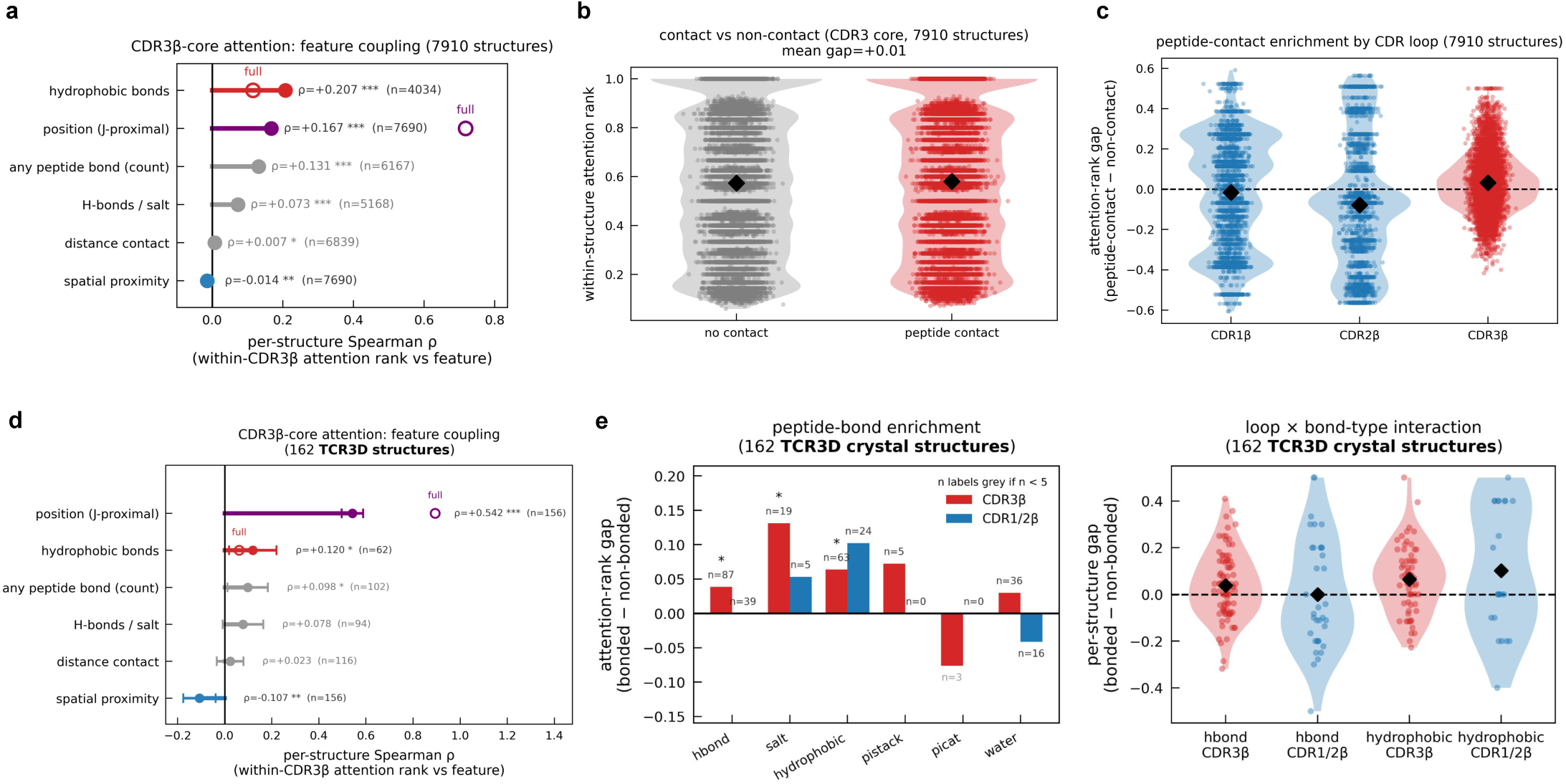
PRISM CDR3β attention tracks peptide-bond chemistry, not contact or proximity; germline CDR1/2β attention is directed away from the peptide. Within-loop attention ranks (scaled to [0,1]) per predicted TCR–pMHC structure (n = 7,910; ipTM ≥ 0.5, CDR3β pLDDT ≥ 70). A residue is a peptide contact if PLIP detects any interaction with the peptide or its heavy-atom min-distance is < 4.5 Å; the trimmed core removes the first and last four CDR3β positions. Points, structures; violins, distributions; diamonds, means. **(a) Per-structure Spearman ρ between CDR3β attention rank and each feature**; filled, trimmed core; open, full loop. Hydrophobic-bond count is the strongest core correlate (ρ = +0.21), above peptide-bond count (+0.13) and hydrogen-bond/salt count (+0.07); neither proximity (min-distance, ρ = −0.01) nor untyped contact (gap +0.01) predicts attention. Position falls from ρ = +0.72 (full loop) to +0.17 (core), identifying it as attention on the fixed J-anchor. All rungs are individually significant, so effect size, not P, is interpretive. Wilcoxon signed-rank test against zero; ***P < 0.001, *P < 0.05. **(b) CDR3β-core attention rank for peptide-contacting versus non-contacting residues**: near-identical (mean rank 0.57 vs. 0.58; gap +0.01). **(c) Per-structure contact gap for CDR1, CDR2 and CDR3 as complete loops.** CDR1 and CDR2 are negative (≈ −0.02, −0.09); the CDR3β gap is marginally positive (+0.03), far below its bond-type enrichments in Fig. 4c. Only the TCR–peptide interface was annotated; germline orientation toward the MHC is not tested here. **(d) Per-structure Spearman ρ between CDR3β attention rank and each feature in 162 TCR3d crystal structures;** filled, trimmed core; open, full loop. Position is the strongest correlate at both resolutions (ρ = +0.89, full loop; ρ = +0.54, core), consistent with a J-gene-anchored positional bias rather than a structural contact signal — position does not track peptide proximity (min-distance ρ = −0.07 to −0.11) or contact frequency in this structure set. Among biochemical features, hydrophobic-bond count is the strongest core correlate (ρ = +0.12), above peptide-bond count (+0.10) and hydrogen-bond/salt count (+0.08); untyped contact (gap = +0.02) does not reach significance. All other rungs are individually significant, so effect size, not P, is interpretive. Wilcoxon signed-rank test against zero; ***P < 0.001, *P < 0.05. **(e) Attention enrichment on peptide-bonding residues, computed from 162 TCR3D crystal structures.** Bonds are specific TCR–peptide interactions identified by PLIP (hydrogen bond, salt bridge, hydrophobic, π-stacking, π-cation), independent of spatial proximity. Within each structure, loop residues were ranked by attention weight (scaled to [0,1]); the attention-rank gap is the mean rank of residues forming a given bond to the peptide minus the mean rank of residues not forming that bond (positive, attention favors bond-forming residues). Left, mean gap per bond type for CDR3β (red) and CDR1/2β (blue); n, structures containing ≥1 bond of that type (grey where n < 5). Right, per-structure gap distributions for the two best-powered bond types (hydrogen bond, hydrophobic), split by loop, where points are structures and diamonds are means. CDR3β attention is significantly enriched on salt bridges (gap = +0.13, P = 0.009), hydrophobic bonds (+0.06, P = 0.002), and weakly on hydrogen bonds (+0.04, P = 0.021); CDR1/2β shows a comparable or larger hydrophobic-bond enrichment (+0.10, n.s.) but no hydrogen-bond enrichment (≈0, n.s.), so the two loops do not show a consistent sign reversal in this structure set. Wilcoxon signed-rank test against zero; ***P < 0.001, *P < 0.05.

